# DNA binding and lesion recognition by the bacterial interstrand DNA crosslink glycosylase AlkX

**DOI:** 10.64898/2025.12.11.693820

**Authors:** Yujuan Cai, Dillon E. Kunkle, Marcel D. Edinbugh, Eric P. Skaar, Brandt F. Eichman

**Affiliations:** Department of Biological Sciences, Vanderbilt University, Nashville, Tennessee 37232; Department of Pathology, Microbiology, and Immunology, Vanderbilt University Medical Center, Nashville, Tennessee 37232; Vanderbilt Institute for Infection, Immunology, and Inflammation, Vanderbilt University Medical Center, Nashville, Tennessee 37232; Department of Biology, Fisk University, Nashville, Tennessee 37208; Department of Biochemistry, Vanderbilt University, Nashville, Tennessee 37232

**Keywords:** Bacterial pathogenesis, DNA repair, host-pathogen interactions

## Abstract

Interstrand DNA crosslinks (ICLs) are a highly toxic form of DNA damage. ICL repair in both eukaryotes and bacteria involves unhooking of the two strands by specialized DNA glycosylases. We recently established that the human pathogen *Acinetobacter baumannii* contains an ICL glycosylase (AlkX) that facilitates pathogenesis and protects the bacteria from DNA damage and acid stress. However, the physical basis for glycosylase-catalyzed ICL unhooking is unknown. Here, we describe a crystal structure of AlkX bound to DNA representing a product of the ICL unhooking reaction. Mutational analysis of ICL unhooking *in vitro* and *A. baumannii* sensitivity to the crosslinking agent mechlorethamine enabled identification of several AlkX motifs critical for ICL repair. We also found that a genetic variant from an antibiotic-resistant strain of the human pathogen *Salmonella enterica* significantly reduced AlkX activity *in vitro* and increased *A. baumannii* sensitivity to DNA crosslinking. This work provides a structural basis for how bacterial ICL glycosylases recognize and repair DNA adducts and contributes additional evidence that ICL repair is important for fitness of human pathogens.

## Introduction

Interstrand DNA crosslinks (ICLs) are a particularly toxic form of DNA damage that covalently tether opposing DNA strands, thus impeding crucial cellular processes such as replication and transcription (Clauson *et al*, 2013; Noll *et al*, 2006; Osawa *et al*, 2011; Schärer, 2005). ICLs are formed from a variety of endogenous cellular metabolites and environmental toxins (Housh *et al*, 2021; Noll *et al*., 2006). For example, reactive aldehydes generated from lipid oxidization or alcohol metabolism generate ICLs between opposing guanines (Niedernhofer *et al*, 2003; Sonohara *et al*, 2019; Voulgaridou *et al*, 2011). Similarly, abasic (apurinic/apyrimidinic, AP) sites formed by spontaneous depurination or DNA base excision repair can form ICLs by reacting with exocyclic amino groups on the opposite strand (Amidon & Eichman, 2020; Dutta *et al*, 2007; Thompson & Cortez, 2020). Microbes and plants also produce crosslinking secondary metabolites (e.g., azinomycin, psoralen, and mitomycin C) (Foulke-Abel *et al*, 2011; Gates, 1999; Hearst, 1989; Semlow & Walter, 2021). Because of their toxicity, crosslinking agents such as mitomycin C and cisplatin are used as antitumor drugs (Rajski & Williams, 1998; Rycenga & Long, 2018).

ICL repair in eukaryotes and prokaryotes involves unhooking the two strands by one of several hydrolytic mechanisms (Bellani *et al*, 2024; Hodskinson *et al*, 2020; McVey, 2010; Semlow & Walter, 2021). One unhooking pathway involves DNA glycosylase cleavage of the *N*-glycosidic bond of one of the crosslinked nucleotides, which produces an abasic site on one strand and a monoadduct on the other (**Fig. 1A**). The abasic site is either replaced by an undamaged nucleotide via the base excision repair (BER) pathway or transiently protected during DNA replication by the HMCES/YedK pathway (Gohil *et al*, 2023; Krokan & Bjørås, 2013; Mohni *et al*, 2019; Semlow *et al*, 2022; Thompson *et al*, 2019; Thompson & Cortez, 2020). Bacterial ICL glycosylases, which belong to the HTH_42 (PF06224) superfamily of proteins, are prevalent in antibiotic-producing and pathogenic strains (Bradley *et al*, 2022; Bradley *et al*, 2020; Mullins *et al*, 2017b; Wang *et al*, 2016). In *Streptomyces sahachiroi*, the DNA glycosylase AlkZ unhooks ICLs derived from the bifunctional alkylating agent azinomycin B (AZB) (Bradley *et al*., 2020; Wang *et al*., 2016). The *alkZ* gene is embedded within the AZB biosynthetic gene cluster and serves as a self-resistance mechanism against AZB toxicity (Chen *et al*, 2022b; Wang *et al*., 2016). The crystal structure of AlkZ revealed a novel architecture in which three winged helix-turn-helix (WH) motifs scaffold a concave, positively charged DNA-binding surface that contains a catalytic QxQ motif (Mullins *et al*., 2017b). In *Escherichia coli*, the HTH_42 protein YcaQ hydrolyzes N7-alkylguanosine adducts, including N7-methylguanosine (7mG) monoadducts and ICLs derived from the nitrogen mustard (NM) mechlorethamine (Bradley *et al*., 2020) (**Fig. 1A**). Genetic analysis showed that YcaQ initiates a secondary ICL repair pathway distinct from the known nucleotide excision ICL repair pathway (Bradley *et al*., 2020).

**Figure 1.**
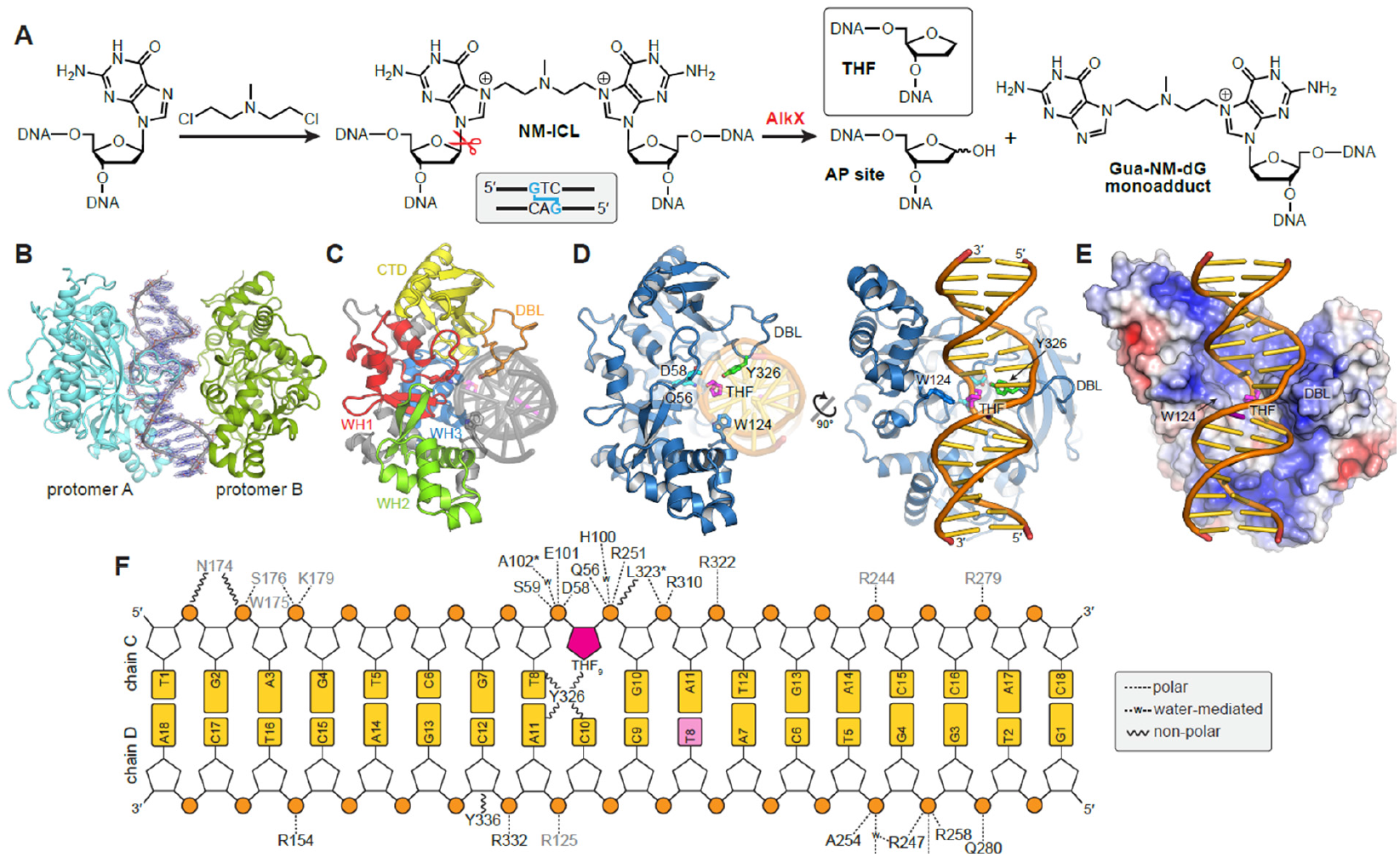
Crystal structure of AlkX bound to DNA. **A**. NM-ICL formation and hydrolysis. The THF abasic site analog used for structure determination is shown as an inset above. **B**. TfuAlkX-DNA asymmetric unit, with 2Fo-Fc electron density contoured to 2σ superimposed on the DNA. **C**. DNA-bound protomer A, colored by domain. **D**. Orthogonal views of TfuAlkX (blue) bound to THF-DNA (orange/gold). The THF moiety is colored magenta. Active site residues Q56 and D58 are cyan and Y326 within the DNA-binding loop is green. **E**. Electrostatic potential surface of TfuAlkX (blue, positive; red, negative), in the same view as the right-hand image in panel C. **F**. Protein-DNA interactions. Hydrogen bonds and polar interactions are shown as dashed lines. Non-polar Van der Waals interactions within 4 Å are shown in wavy lines. Protein residues are colored according to promotor (A, black; B, grey). Asterisks denote main chain contacts.

AlkZ and YcaQ represent two distinct subgroups within the HTH_42 superfamily (Bradley *et al*., 2022). Proteins within the AlkZ-like (AZL) subgroup are enriched in BGCs of antibiotic-producing bacteria and are highly specific for DNA adducts formed by their cognate natural product, thus serving an anti-toxin role. Outside of their conserved Q/HxQ catalytic motifs, AZLs have relatively low sequence similarity, consistent with their varied substrate specificities (Bradley *et al*., 2022; Chen *et al*, 2022a; Mullins *et al*., 2017b). In contrast, the YcaQ-like (YQL) proteins are not found in BGCs, are prevalent in pathogenic bacteria, and share high sequence conservation, including a QxD catalytic motif (Bradley *et al*., 2022; Bradley *et al*., 2020). Thus, whereas AZLs have evolved specificity for a particular secondary metabolite in an antibiotic producer, YQLs can unhook a variety of ICLs and thus likely provide a more general defense against endogenous and exogenous genotoxins. Although the crystal structure of AlkZ has been determined, the structural features of the YQL subfamily are unknown.

We recently characterized an *Acinetobacter baumannii* YQL ortholog, which we named AlkX, as an ICL DNA glycosylase that protects the bacteria against mechlorethamine toxicity (Kunkle *et al*, 2024). *A. baumannii* is an antibiotic-resistant Gram-negative nosocomial pathogen that was the second leading cause of global antibiotic-resistant attributable deaths in 2021 (Collaborators, 2024). *A. baumannii* strains that have evolved resistance to last-resort antibiotics are commonly isolated from patients, leading the WHO to identify *A. baumannii* as a critical priority pathogen, indicating the need for the development of new antibacterials to treat these drug-resistant infections (Sati *et al*, 2025). AlkX promotes *A. baumannii* colonization of the lungs and dissemination to distal tissues during pneumonia. In addition to protecting cells against ICL agents, *alkX* is induced by, and protects against, acid stress, a condition the bacteria encounter while interacting with immune cells in the host (Kunkle *et al*., 2024).

Despite the importance of AlkX for *A. baumannii* fitness and of AlkX, and other YQL DNA glycosylases for ICL repair in prokaryotes, the mechanism by which they unhook ICLs is poorly understood. To fill this gap in knowledge, we determined a crystal structure of AlkX bound to DNA containing an abasic site, representing the product of the ICL unhooking reaction. DNA binding by the HTH_42 architecture places the catalytic QxD motif at the lesion using two DNA binding motifs that clamp the damage site from both major and minor grooves. Mutational analysis of these motifs in AlkX verified their importance for substrate binding, ICL unhooking, and *A. baumannii* fitness. We also identified a critical structural feature of the active site that would be perturbed in a putative AlkX variant found in an antibiotic-resistant strain of another human pathogen, *Salmonella enterica* (Jones-Dias *et al*, 2016), providing additional evidence for the importance of ICL unhooking activity by this family of proteins in pathogenic bacteria. Finally, comparison of the AlkX and AlkZ structures provides a basis for the substrate differences in the YQL and AZL subfamilies. Together, these results provide molecular insight into recognition and repair of ICLs by DNA glycosylases important for viability and pathogenesis in bacteria.

## Results and Discussion

### Crystal structure of AlkX bound to DNA

After screening several YQL/AlkX orthologs and oligonucleotides, we obtained diffraction quality crystals of the *Thermobifida fusca* enzyme (TfuAlkX) bound to DNA. TfuAlkX unhooks NM-ICLs and excises 7mG monoadducts (Bradley *et al*., 2022) and shares 30% sequence identity and 47% similarity to *A. baumannii* (Aba) AlkX and 34% identity and 50% similarity to *E. coli* YcaQ (**Fig. S1**). TfuAlkX was crystallized with an 18-mer DNA oligonucleotide containing a centrally located tetrahydrofuran (THF) abasic site analog, mimicking the product of the DNA glycosylase reaction (**Fig. 1A**). The structure was determined to 2.6 Å resolution by single-wavelength anomalous dispersion (SAD) from selenomethionine-substituted protein and refined to a crystallographic residual (*R*-factor) of 20.6% (R_free_=25.8%) (**Fig. S2, Table S1**). The asymmetric unit contained two protomers and one DNA duplex, with only one protomer engaged with the DNA (**Fig. 1B**).

TfuAlkX adopts a 3xWH/β-barrel architecture similar to that of *Streptomyces sahachiroi* AlkZ (**Fig. 1C, Fig. S3**). The RMSD between TfuAlkX and AlkZ is 3.99 Å for 269 C_α_ atoms, highlighting the conservation of the HTH_42 fold. Consistent with our previous prediction for AlkZ (Mullins *et al*., 2017b), TfuAlkX binds the DNA against its concave, positively-charged surface, with the THF moiety close to the catalytic QxD motif (**Fig. 1D,E, Fig. S2**). Although N7-methylguanine (7mGua) nucleobase was included in the crystallization condition to stabilize the enzyme-product complex (Mullins *et al*, 2015), it was not evident in the electron density. Instead, a molecule of ethylene glycol from the crystallization condition was observed to fill the nucleobase void beside the THF abasic site (**Fig. S2**), suggesting that AlkX does not remain bound to the excised nucleobase product. The THF (lesion) site is clamped between an extensive DNA-binding loop (DBL) from the β-sheet subdomain in the minor groove and a tryptophan side chain (W124) from the region between WH1 and WH2 in the major groove (**Fig. 1D-F**). Thus, the resolution and quality of the electron density in the structure reveal, for the first time, several critical interactions between a bacterial ICL glycosylase and damaged DNA.

### The DNA binding loop is important for AlkX function

Perhaps the most striking feature of the TfuAlkX structure is the DBL protruding into the minor groove at the lesion site (**Fig. 1D-E**). The DBL is located within the β-sheet subdomain and contains 18 residues (F319-Y336), nine of which (E324-R332) adopt a cradle for the phosphoribose backbone of the strand opposite the THF (**Fig. 2A**). At the tip of the loop, a conserved tyrosine (Y326) partially fills the abasic site void and forms van der Waals contacts with the THF and the nucleobases around the THF. The DBL is in a similar position as the β11/12 motif of AlkZ that we previously showed to be essential for glycosylase activity (**Fig. S3**) (Mullins *et al*, 2014). To determine the importance of the DBL in AlkX function we generated mutants within the corresponding region in AbaAlkX and examined NM-ICL unhooking activity *in vitro* (**Fig. 2B, Fig. S4**). In this assay, AlkX is incubated with a Cy5-labeled NM-ICL substrate, which generates a guanine-NM-dG monoadduct (MA) strand and an AP strand (**Fig. 1A**) that can be separated by electrophoresis after alkali nicking of the AP strand (Bradley *et al*., 2020). ICL unhooking was severely curtailed in a “Δloop” mutant in which AbaAlkX loop residues E298-R306 (corresponding to E324-R332 in TfuAlkX) were replaced with GSSG (**Fig. 2B**). Similarly, ICL unhooking was reduced by alanine substitution of the tyrosine side chain (Y300A). We verified that the reduction in activity by the mutants was not the result of reduced protein thermostability (**Fig. S5**). Phenylalanine substitution of Y300 had no effect on ICL unhooking, indicating that the hydroxyl group is not important for catalysis. The corresponding mutants in TfuAlkX—Δloop (E324-R332>GSSG), Y326A, and Y326F—showed the same effect as those in AbaAlkX (**Fig. S6**).

**Figure 2.**
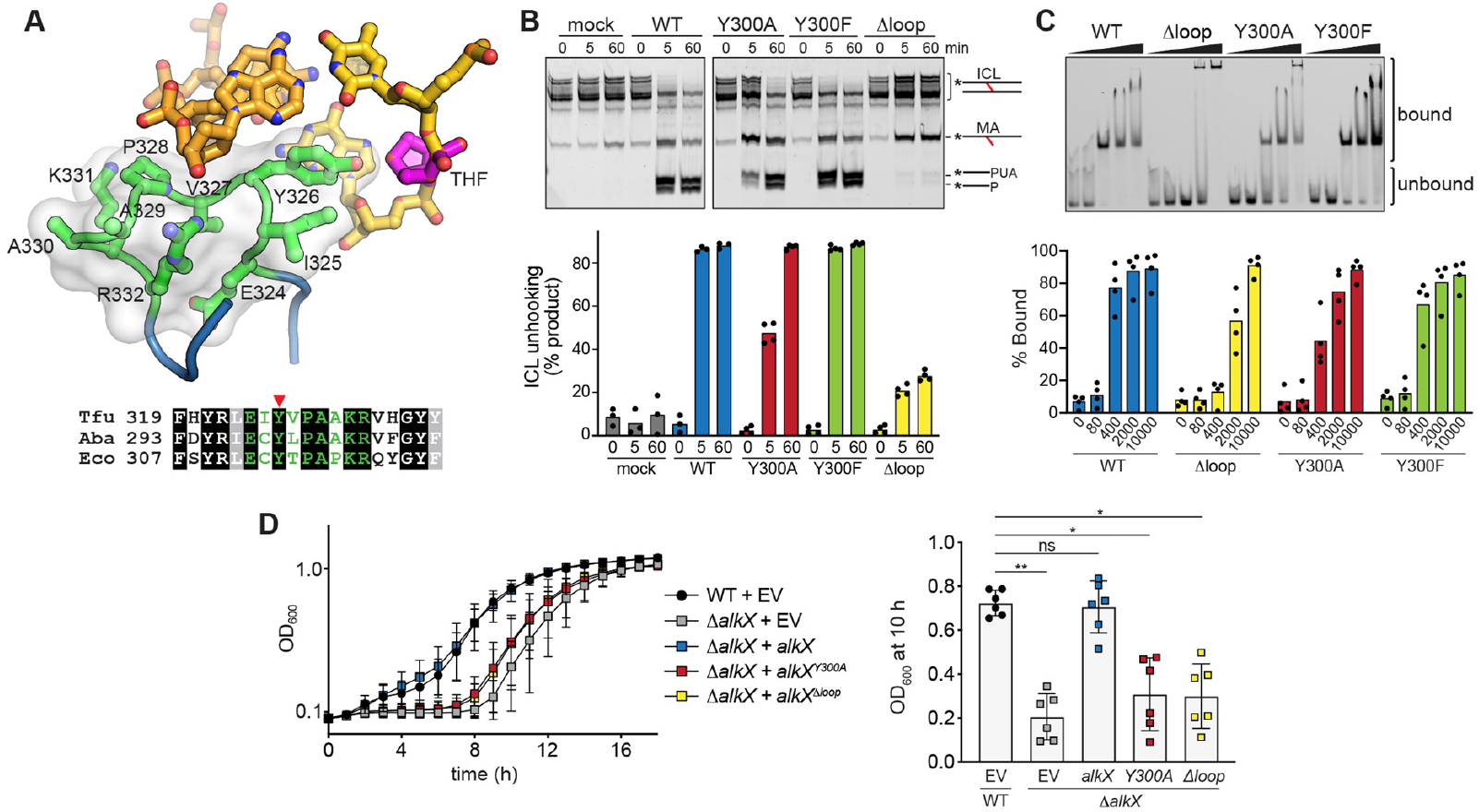
The DNA-binding loop (DBL) is important for AlkX function. **A**. Structure of the DBL (green) in contact with the DNA minor groove at the lesion site (magenta THF). The solvent accessible surface of the DBL is shown in transparent white. The sequence alignment of the DBL between AlkX/YQL orthologs from *Thermobifida fusca* (Tfu), *Acinetobacter baumannii* (Aba), and *Escherichia coli* (Eco) is shown below. The conserved tyrosine is highlighted with a red triangle. **B**. NM-ICL unhooking by AbaAlkX wild-type (WT) and mutants. A representative denaturing polyacrylamide gel shows conversion of NM-ICL substrate to a monoadduct (MA) strand and alkali-cleaved AP-strand products. PUA, 3′-phospho-α,β-unsaturated aldehyde; P, 3′-phosphate; mock, no enzyme. Quantification shows the mean of 3-4 independent experiments. **C**. Electrophoretic mobility shift assay (EMSA) of AlkX mutants binding to 2F-NM_8_-ICL DNA (Fig. S2). Quantification of four replicate experiments is shown below. **D**. Growth curves of *A. baumannii* WT and Δ*alkX* strains harboring either empty vector (EV) or indicated *alkX* expression vectors in the presence of 33 µM mechlorethamine. Data represent the mean ± SD of six biological replicates performed in technical triplicate. The comparison of strain viability harboring different mutants is shown on the right with the OD_600_ values at 10 hr. Data represent the mean ± SD, each dot represents an individual biological replicate performed in technical triplicate. *P < 0.5, **P < 0.01, ns (not significant) as determined by Dunn’s multiple comparisons test.

We also probed the effect of the mutations on AbaAlkX binding to DNA containing a stable NM-ICL constructed of an 8-atom NM analog, which we previously showed to be a substrate for *E. coli* YcaQ (Bradley *et al*., 2020), and C2′-fluorinated guanines to prevent glycosylase cleavage (**Fig. 2C, Fig. S4**). Similar to the ICL unhooking results, DNA binding was severely curtailed by the Δloop mutant, reduced by Y300A, and unaffected by Y300F, indicating that the tyrosine contributes sterically to DNA binding by the DBL. Consistent with the importance of the DBL in DNA binding, electron density for the DBL was not observed in the protomer not engaged with the DNA, suggesting that this motif is disordered in the absence of DNA.

Given the effect of Δloop and Y300A mutants on ICL unhooking and DNA binding, we probed the effect of these mutations *in vivo* by measuring the ability of them to complement an *A. baumannii ΔalkX* mutant strain grown in the presence of the DNA crosslinker mechlorethamine. Consistent with the *in vitro* results, the DBL mutants increased the sensitivity of *A. baumannii* DNA crosslinking, as monitored by cell growth (**Fig. 2D, Fig. S4**). The *ΔalkX* strain showed greater sensitivity compared to WT as judged by delayed growth. This sensitivity was complemented by expression of wild-type *alkX*. However, expression of either *Y300A* or *Δloop alkX* mutant alleles failed to complement the loss of *alkX*. Thus, cells harboring these mutants exhibit greater mechlorethamine sensitivity. Collectively, these findings have uncovered a previously unknown region of the protein that is required for ICL repair. Based on the extremely high conservation of the DBL across YQL/AlkX proteins, it stands to reason that these proteins will process ICLs in the same manner.

### A genetic variant in *S. enterica* impairs ICL unhooking

The glutamine and aspartate residues within the conserved catalytic QxD motif are essential for base excision in *E. coli* YcaQ and AbaAlkX (Bradley *et al*., 2020; Kunkle *et al*., 2024). We postulate that one or both of these residues stabilize the position of the catalytic water for attack of the lesion C1′ (Mullins *et al*, 2019; Mullins *et al*., 2017b). In the TfuAlkX crystal structure, the Q56 and D58 side chains of the QxD motif form hydrogen bonds with the phosphate groups 5′ and 3′ to the THF nucleotide (**Fig. 3A**). As a result, neither side chain is close enough to the C1′ carbon to catalyze hydrolysis, and thus the structure suggests that the enzyme and/or DNA undergo a small conformational change after cleavage of the N-glycosidic bond, as observed in other alkylpurine DNA glycosylases (Metz *et al*, 2007). The phosphate interactions impart a deformation in the THF backbone such that the THF ring is rotated 90° relative to the position of the deoxyribose rings in duplex DNA, with the C1′ carbon facing the DNA duplex (**Fig. 3A**). This position, together with the lack of empty space between the THF and the protein, is consistent with a non-base-flipping catalytic mechanism expected for an ICL glycosylase (Mullins *et al*., 2019). Although the structure revealed a glutamate (E101) side chain close to the C1′ carbon, alanine substitution of this residue in TfuAlkX (E101A) did not affect ICL unhooking activity (**Fig. S6**), and thus we conclude that it likely does not participate in catalysis. This was not unexpected since this residue, while conserved in most AlkX/YQL orthologs, is an alanine in AbaAlkX (**Fig. 3B, Fig. S1**).

**Figure 3.**
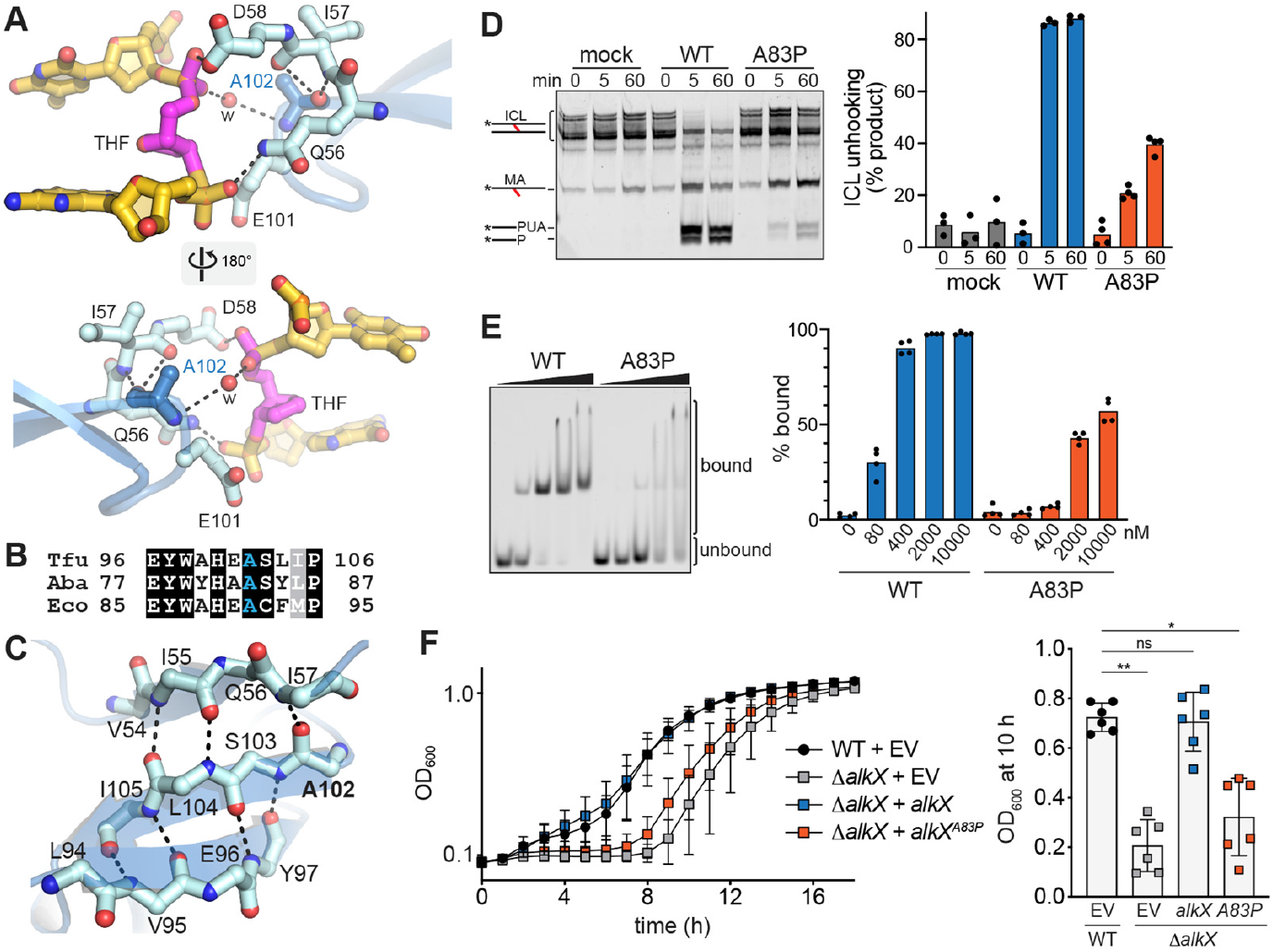
A genetic variant in *S. enterica* impairs ICL unhooking. **A**. Crystal structure of the active site, showing the proximity of A102 main chain to the catalytic QxD motif. Hydrogen bonds are shown as dashed lines and a bridging water is shown as a red sphere. **B**. Sequence alignment of this region in AlkX**/**YQL proteins from *T. fusca* (Tfu), *A. baumannii* (Aba), and *E. coli* (Eco). **C**. A102 main chain is linked to the QxD motif within a β-sheet in the WH1 domain. Only main chain atoms are shown. **D**. Representative gel and quantification of ICL unhooking activity of AbaAlkX A83P. Bars denote the mean from 3-4 replicates. **E**. Representative EMSA of AbaAlkX A83P binding to 2F-NM_8_-ICL, with quantification showing the mean from four independent measurements. **F**. Growth curves of *A. baumannii* WT and *ΔalkX* strains harboring empty vector (EV) or indicated *alkX* expression vectors under in the presence of 33 µM mechlorethamine. Data represent the mean ± SD of six biological replicates performed in technical triplicate. Comparison of strain viability harboring *alkX* WT and A83P is shown on the right with the OD_600_ values at 10 hr. Data represent the mean ± SD, each dot represents an individual biological replicate. *P < 0.5, **P < 0.01, ns (not significant) as determined by Dunn’s multiple comparisons test.

The structure revealed an important element of the active site that likely stabilizes the positions of both the QxD motif and the DNA lesion. At the end of the β-hairpin in WH1 (the “wing”

of the winged-helix), the main chain atoms of residue A102 form a direct hydrogen bond with the main chain of I57 within the QxD (QID) motif and a water-mediated hydrogen bond with the THF phosphate (**Fig. 3A, 3C**). Highly conserved in AlkX/YQL proteins (**Fig. S1**), A102 is also involved in a three-stranded β-sheet and thus is an important structural feature that imparts stability to the active site (**Fig. 3C**). Interestingly, a corresponding alanine in a putative YQL ortholog in *Salmonella enterica* is mutated to a proline in a single nucleotide variant (A89P) found in a tetracycline-resistant strain of *S. enterica* serotype Enteritidis (Jones-Dias *et al*., 2016). *S. enterica* is a dangerous human pathogen that causes gastrointestinal infections related to the consumption of contaminated animal products (Foley *et al*, 2013). Based on the importance of this alanine to the structure of the AlkX active site, the *S. enterica* A89P mutation likely alters the position of the catalytic motif to consequently impair ICL unhooking activity. Consistent with this assertion, we found that proline substitution of the corresponding alanine in AbaAlkX (A83P) dramatically reduced ICL unhooking and ICL-DNA binding *in vitro* (**Fig. 3D-E, Fig. S4**) without destabilizing the thermostability of the protein (**Fig. S5**). *In vivo*, the AbaAlkX A83P mutation sensitized the *ΔalkX A. baumannii* strain to mechlorethamine to the same extent as the DBL and QxD mutations (**Fig. 3F, Fig. S4**) (Kunkle *et al*., 2024). We confirmed that a TfuAlkX A102P mutant also impaired ICL unhooking activity (**Fig. S6**). Thus, the WH1 motif is a conserved feature in AlkX orthologs that stabilizes the catalytic QxD motif for ICL unhooking. Furthermore, our structure of an AlkX/YQL ICL glycosylase from a pathogenic bacterium has provided a possible mechanistic explanation for how a genetic variant found in an antibiotic-resistant strain of *S. enterica* inactivates the glycosylase.

Although the isolated *S. enterica* strain contains an inactivating mutation in its AlkX homologue, the strain maintains virulence. This contrasts with *A. baumannii* AlkX, which is highly conserved in *A. baumannii* clinical isolates, does not contain this inactivating mutation, and aids in colonization of the mouse lung. Together, these findings suggest that the selective pressures on these two YQL proteins differ in these individual organisms. Indeed, although DNA repair pathways are essential to the virulence of several important human pathogens, including *A. baumannii* (Morris *et al*, 2025; Zgur-Bertok, 2013), the inactivation of DNA repair genes, which increases genome instability, has been suggested to be an important mechanism to produce genetic variation and adaptation. Loss of DNA repair genes has been attributed to increased evolution rates, mutator phenotypes, and increased drug resistance in microbes (Billmyre *et al*, 2017; Boyce, 2022; Boyce *et al*, 2017; Oliver *et al*, 2000; Rhodes *et al*, 2017; Steenwyk *et al*, 2019; Zein-Eddine *et al*, 2025). Whether the A83P variant in *S. enterica* AlkX engenders this organism with similar characteristics and contributes to tetracycline resistance remains to be determined.

### Identification of a putative ICL sensor in AlkX

On the opposite side of the DNA duplex from the DBL, W124 penetrates the major groove at the lesion site (**Fig. 1D-E, Fig. 4A**). W124 is located at the N-terminal end of helix αF, in the region connecting WH2 and WH4 (**Fig. S3**). The W124 side chain does not make any DNA contacts in the structure. However, modeling a NM onto the DNA suggests that W124 would contact the crosslink (**Fig. 4A**). Indeed, mutation of W124 to alanine in TfuAlkX reduced ICL unhooking relative to the wild-type protein, although to a lesser extent than that of the DBL Y326A mutant on the other side of the DNA or the A102P mutant in the active site (**Fig. 4B, Fig. S6**). Nonetheless, the position of this residue and its reduction in NM-ICL activity suggests that it plays a role in sensing the crosslink. Consistent with this, this αF region is relatively well conserved among YQL proteins but divergent among AZLs, which have evolved specificities for DNA adducts of a particular natural product (Bradley *et al*., 2022; Bradley *et al*., 2020; Chen *et al*., 2022a). It is not surprising, therefore, that the conformations of the αF regions in the TfuAlkX and *Streptomyces sahachiroi* (Ssa) AlkZ structures are different (**Fig. 4C**). This difference, however, may simply reflect a DNA-bound (AlkX) versus DNA-free (AlkZ) conformation. Interestingly, although the lesion-sensing tryptophan in TfuAlkX is highly conserved among AlkX/YQL proteins, it is a serine in AbaAlkX (**Fig. 4A, Fig. S1**). Thus, it is possible that this motif discriminates among ICLs or minor groove lesions in an organism-specific manner.

**Figure 4.**
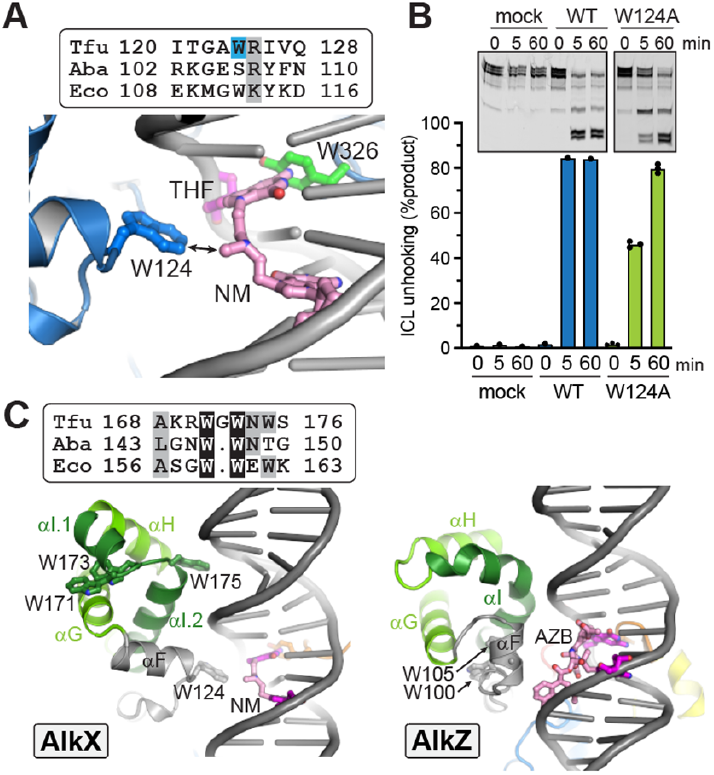
Identification of a putative ICL sensor in AlkX. **A**. Model of TfuAlkX bound to the full product of the NM-ICL unhooking reaction, based on the TfuAlkX/THF-DNA structure. The Gua-NM-dG monoadduct is shown in pink, and the THF from the crystal structure is magenta. A sequence alignment of the αF region in YQLs from *T. fusca* (Tfu), *A. baumannii* (Aba), and *E. coli* (Eco) is shown above. **B**. ICL unhooking activity of TfuAlkX W124A. Data represent the mean from three independent replicates. **C**. Comparison of αF (grey) and WH2 (green) regions of AlkX (left) and AlkZ (right). DNA models of NM- and azinomycin B (AZB)-ICLs are based on the DNA in the TfuAlkX structure. A sequence alignment of the αI region in AlkX orthologs is shown above.

We previously predicted the αI helix within WH2 to serve as a lesion recognition motif in the Streptomyces AZL proteins AlkZ, TxnU2/4, and LldU1/5, based on its high sequence divergence and predicted interaction with the lesion site in models constructed from the AlkZ structure (Chen *et al*., 2022a; Mullins *et al*., 2017b). In contrast, in TfuAlkX, WH2 contacts the DNA one-half turn up the duplex from the αF region (**Fig. 4C**) and therefore does not contact the lesion. Helix αI, which is highly kinked in AlkZ, is separated into two shorter alpha helices and connected with a triple tryptophan loop (WGWNW) in TfuAlkX. W171 and W173 face into the WH2 core while W175 points toward the DNA minor groove (**Fig. 4C, Fig. S3**). Consistent with the structure, substitution of W175 with alanine only modestly reduced ICL unhooking by TfuAlkX, and a W173A mutant had no effect (**Fig. S7**). W171 and W173, but not W175, are conserved in AbaAlkX, and mutation of these residues had no effect on ICL unhooking in AbaAlkX (**Fig. S7**). Thus, we conclude that the WH2 region does not play a role in ICL unhooking in the YQL/AlkX subfamily. It remains to be determined if this region serves as an ICL recognition motif in the AZLs. Modeling AlkZ against an AZB-ICL using the DNA from the TfuAlkX structure does not place helix αI in proximity to the crosslink. However, the current structure represents a relaxed (non-crosslinked) DNA conformation, and thus it is difficult to approximate the proximity of the αF and αI helices to the lesion in a true ICL substrate. Future investigation into this region is needed to discern its effect on the substrate specificities of the AZL glycosylases.

A putative ICL sensor in AlkX that differs between TfuAlkX and AbaAlkX raises the possibility that AlkX/YQL proteins have evolved to serve different DNA repair processes in individual species. This hypothesis is supported by the observation that *E. coli ycaQ* transcription is under the regulation of a σ^70^-dependent promoter, and this architecture is conserved in *Salmonella enterica*, suggesting that the YQL genes in these organisms are constitutively expressed (Bradley *et al*., 2020). However, *alkX* expression in *A. baumannii* is induced in response to mechlorethamine treatment, and treatment of *P. aeruginosa* with a quorum sensing inhibitor induces expression of its *alkX* homologue (Tan *et al*, 2014). Collectively, these findings suggest the possibility that distinct organisms have evolved alterations in YQL protein structures as well as species-specific regulatory mechanisms to control expression of YQL proteins that serve specific functions in distinct niches.

### Conclusion and outlook

To our knowledge, the present work reports the first structural information for how a bacterial ICL glycosylase engages DNA and achieves specificity for an ICL. Our AlkX-DNA structure revealed several conserved motifs that we found to be important for ICL unhooking and recognition, mutation of which led to increases in *A. baumannii* sensitivity to DNA crosslinking. Given the widespread distribution of AlkX/YQL ICL glycosylases in pathogenic bacteria, including

*A. baumannii, Salmonella enterica, Listeria monocytogenes*, and *Pseudomonas aeruginosa*, it will be important to determine the role of these enzymes in pathogenesis and to define their endogenous substrates.

While DNA glycosylases traditionally have been regarded as a means of repairing nucleobases containing small modifications from oxidation, deamination, and alkylation as a component of the base excision repair (BER) pathway, they are now established as a means of repairing bulky adducts and ICLs in both prokaryotes and eukaryotes (Baiken *et al*, 2020; Bradley *et al*., 2022; Bradley *et al*., 2020; Kothandapani & Patrick, 2013; Mullins *et al*., 2019; Mullins *et al*, 2017a; Mullins *et al*., 2015; Semlow *et al*, 2016; Wang *et al*., 2016; Xu *et al*, 2012). The bacterial ICL glycosylases characterized to date (AlkZ, AlkX/YcaQ) seem to be specific for compounds that link guanines through the N7 nitrogen (Bradley *et al*., 2020; Gates *et al*, 2004; Mullins *et al*., 2017b; Wang *et al*., 2016), and related glycosylases in the HTH_42 enzyme family recognized N7-alkylated monoadducts (Bradley *et al*., 2022; Chen *et al*., 2022a). In contrast, the eukaryotic DNA glycosylase NEIL3 (Endonuclease VIII like 3) repairs AP- and psoralen-ICLs as an alternative to the Fanconi Anemia (FA) pathway and to process double-strand breaks in FA-dependent repair of mitomycin C and cisplatin-ICLs (Li *et al*, 2020; Li *et al*, 2022; Oswalt & Eichman, 2024; Semlow *et al*., 2016). Unlike the bacterial ICL glycosylases, NEIL cannot unhook ICLs from double-stranded DNA; it requires a double/single-stranded (splayed arm) junction that would be present at replication forks, consistent with its cellular recruitment to ICLs present at converged forks (Imani Nejad *et al*, 2020; Semlow *et al*., 2016; Wu *et al*, 2019). Moreover, the NEIL3 glycosylase domain adopts an entirely different fold than AlkX and does not contain the same DNA-penetrating motifs that we identified for AlkX (Huskova *et al*, 2022; Liu *et al*, 2013). Thus, it remains to be determined how NEIL3 achieves specificity for ICLs at branched DNA structures, which may require domains outside of its glycosylase domain (Huskova *et al*., 2022; Imani Nejad *et al*., 2020; Rodriguez *et al*, 2020).

## Methods

### Reagents

All chemicals were purchased from Sigma-Aldrich unless otherwise noted. *N1*,*N2*-bis(2-chloroethyl)-*N1*,*N2*-dimethylethane-1,2-diamine (NM_8_) used in the preparation of EMSA substrates was synthesized as previously described (Bradley *et al*., 2020). All plasmids and primers used are listed in **Table S2**. DNA oligonucleotides were purchased from Integrated DNA Technologies.

### Cloning

*AlkXA83P* and *Y300A* mutant allele expression constructs were generated by site-directed mutagenesis as follows. The pWH1266-*alkX* vector was amplified in a PCR reaction with primers encoding the indicated point mutants. The resulting PCR product was DpnI digested and transformed into *E. coli*. The *alkXΔloop* mutant allele expression construct was generated by amplifying the *alkX* promoter and the *alkX* allele in which E298-R306 was replaced with GSSG from a gBlock (IDT). The resulting fragment was cloned into a SalI and BamHI digested pWH1266 vector using HiFi Assembly. Plasmids harboring the desired mutations were screened by Sanger sequencing.

### Preparation of ICL-DNA substrates

NM_5_-ICL and 2F-NM_8_-ICL DNA substrates used in ICL unhooking and DNA binding experiments were prepared as previously described (Bradley *et al*., 2020). For the NM_5_-ICL substrate, 26-bp oligodeoxynucleotides Cy5-d(TTTATTTTTATTTGACTTTTATTTTT) and FAM-d(AAAAATAAAAGTCAAATAAAAATAAA) were annealed at 200 μM in 40 mM sodium cacodylate (pH 7.0). The annealed duplex was incubated with 3 equivalents of mechlorethamine•HCl for 3 hr at 37°C in the dark. The reaction mix was desalted using MicroSpin G-25 columns (Cytiva) and gel purified with 15% precast TBE-Urea gels (ThermoFisher). Samples were mixed with loading buffer (5 mM EDTA (pH 8.0), 80% (wt/vol) formamide, 0.5mg/ml Orange G) prior to loading. The gel was pre-run for 1 hr at 200 V and run with samples for 1 hr at 200V. The NM_5_-ICL DNA band was excised and transferred to a 1-kDa MWCO dialysis tube (SpectrumLab), followed by electrophoresis at 100V for 1 hr. After dialysis in TE buffer overnight, the substrate was concentrated to 2 μM, aliquoted, and stored at -80°C. 2F-NM_8_-ICL DNA substrates were prepared using the same procedure, with NM_8_ compound and oligodeoxynucleotides Cy5-TTTATTTTTATTTG^F^ACTTTTATTTTT and FAM-AAAAATAAAAG^F^TCAAATAAAAATAAA, where G^F^ is a ribo-fluoro-C2′ deoxyguanosine

### Protein purification

Wild-type TfuAlkX and AbaAlkX genes were synthesized and cloned into a pBG102 vector (Vanderbilt University Center for Structural Biology) with codon optimization by GenScript. Mutants were generated by PCR using CloneAmp HiFi PCR premix (Takara) and primers containing the mutations (**Table S2**). PCR products were digested with DpnI, gel purified with QIAquick gel extraction kit (QIAGEN), and assembled using HiFi Assembly enzyme (NEB) prior to transformation into DH5α competent cells. All vectors were confirmed by Sanger or Oxford Nanopore sequencing.

AbaAlkX and TfuAlkX proteins were expressed in *E. coli* Tuner (DE3) cells grown in LB media containing 30 µg/mL kanamycin. Expression was induced with 0.1 mM IPTG (isopropyl-β-D-thiogalactopyranoside) at an OD_600_ of 0.8. Selenomethionine-derivatized (SeMet) TfuAlkX was overexpressed in *E. coli* Tuner (DE3) cells with 0.1 mM IPTG induction in M9 media containing 0.1 g/L lysine, 0.1 g/L phenylalanine, 0.1 g/L threonine, 0.05 g/L leucine, 0.05 g/L isoleucine, 0.05 g/L valine, and 0.05 g/L selenomethionine. After growing for 16 hr at 16 °C, cells were harvested, homogenized in lysis buffer (50 mM Tris (pH 8.0), 500 mM NaCl, 1 mM Tris (2-carboxyethyl) phosphine (TCEP), 1 mM phenylmethylsulfonyl fluoride (PMSF), 25 mM imidazole, 10% glycerol) and lysed by sonication. Cell debris was removed by centrifugation at 21,000 rpm for 30 min at 4 °C. The supernatant was loaded onto an Ni-NTA affinity column (Cytiva) and the His_6_-tagged protein was eluted with buffer B (50 mM Tris (pH 8.0), 500 mM NaCl, 250 mM imidazole, 10% glycerol). Protein fractions were pooled and supplemented with 0.1 mM EDTA and 1 mM TCEP before incubation with 1 mg Rhinovirus 3C (PreScission) protease at 4 °C overnight. The cleaved protein was diluted 5-fold with buffer C (50 mM Tris (pH 8.0), 10% glycerol, 0.1 mM EDTA, 1 mM TCEP) and purified on a heparin sepharose column (Cytiva) with a 0–1 M NaCl/buffer C linear gradient. Fractions were pooled and repassed over the Ni-NTA affinity column. The flow-through was concentrated using a 10-kDa MWCO Amicon Ultra filter (Millipore). The concentrated protein was then passed over a Superdex 200 column (Cytiva) in 25 mM Tris (pH 8.0), 150 mM NaCl, 5% glycerol, 0.1 mM EDTA, and 1 mM TCEP. AbaAlkX and TfuAlkX were concentrated to 5 mg/mL and 0.6 mg/mL, respectively, flash-frozen in liquid nitrogen, and stored at -80 °C.

### Crystallization, X-ray data collection, and structure determination

THF-DNA was prepared by annealing d(TGAGTCGT(THF)GATGACCAC) and d(GTGGTCATCCACGACTCA) at 500 μM in the presence of 2 equivalents of N7-methylguanine in 10 mM MES (pH 6.5) and 40 mM NaCl. For crystallization, SeMet-TfuAlkX was incubated with 1.2 equivalents of THF-DNA on ice for 20 min and concentrated to 1.5 mg/ml (protein concentration) using a 30-kDa Amicon MWCO Ultra Centrifugal Filter. Block shaped crystals were observed in hanging drop vapor diffusion plates after 3 days in 0.1 M MES/imidazole (pH 6.5), 10% PEG4000, 10% glycerol, 0.03 M sodium fluoride, 0.03 M sodium bromide, and 0.03 M sodium iodide. Crystals were transferred to a cryoprotectant solution consisting of mother liquor supplemented with 15-20% (v/v) glycerol and flash-frozen in liquid nitrogen for data collection.

X-ray diffraction data (**Table S1**) were collected at the European Synchrotron Radiation Facility (ESRF) beamline ID30A (λ = 0.96546 Å). The datasets were processed using autoPROC (Vonrhein *et al*, 2011) and STARANISO (Vonrhein *et al*, 2018). Phases were determined by single wavelength anomalous dispersion (SAD). Thirteen selenium sites were identified in the asymmetric unit by HySS in the PHENIX package (Adams *et al*, 2010) and used for model building through AutoSol and Autobuild. Iterative cycles of model rebuilding and refinement were carried out using COOT (Emsley *et al*, 2010) and PHENIX (Adams *et al*., 2010). MolProbity was used to access the overall quality of the structural models (Davis *et al*, 2007). Structure figures were made using PyMol 3.0 (Schrödinger, LLC). The RMSD was calculated using TM align in the RCSB online tool (Bittrich *et al*, 2024). All structural biology software was curated by SBGrid (Morin *et al*, 2013).

### Base excision assay

Glycosylase reactions were carried out at room temperature with 1 µM enzyme and 50 nM DNA substrate in glycosylase buffer (50 mM HEPES pH 7.5, 100 mM KCl, 10 mM EDTA, and 10% glycerol). At various time points, 4 µL aliquots were taken and added to 1 µL of 1 M NaOH. The samples were heated at 55°C for 2 min, followed by addition of 5 µL of denaturing loading buffer (5 mM EDTA (pH 8.0), 80% formamide, 1 mg/ml blue dextran) and heating at 55°C for 5 min to avoid spontaneous depurination at high temperature. All samples were electrophoresed on a 20% acrylamide/8 M urea denaturing gel at 40 W for 2 hr in 0.5× TBE buffer. Gels were imaged on an Amersham Typhoon RGB (Cytiva) using the Cy5 (655-685 nm) channel. Bands were quantified with ImageQuant (Cytiva). ICL unhooking (% product) was calculated as (MA + β/δ-elim) / (ICL + MA + β/δ-elim), where ICL, MA, and β/δ-elim are the amounts of ICL substrate, monoadduct product, and AP β/δ-elimination products, respectively.

### Electrophoretic mobility shift assay (EMSA)

DNA binding reactions were carried out by incubating 4 µL protein at varying concentrations with 4 µL 20 nM 2F-NM_8_-ICL DNA substrate at room temperature for 20 min in binding buffer (25 mM Tris (pH 7.5), 100 mM NaCl, 5% glycerol, 0.1 mM EDTA, 1 mM TCEP). Each sample was mixed with 4 µL of 75% glycerol and then loaded onto a 5% native TBE gel. The gel was pre-run at 200 V for 1 hr and run with samples at 200 V for 1 hr. Gels were imaged on a Typhoon laser-scanner platform (Cytiva) using the Cy5 channel and bands quantified with ImageQuant (Cytiva).

### Bacterial growth curves

Overnight cultures of WT and Δ*alkX A. baumannii* strains harboring the indicated plasmids were started from single colonies in LB broth, with 75 µg/mL carbenicillin. The following day overnight cultures were back-diluted into fresh LB media with carbenicillin 1:50, and allowed to grow with shaking at 37 °C for 1 hr. Back-diluted cultures were used to inoculate the wells of 96-well microtiter plates 1:100 with carbenicillin ± 33 µM mechlorethamine-HCL. Plates were cultured with continuous shaking at 37 °C for 18 hr and OD_600_ was recorded every 60 min.

## Data Availability

Structure factors and corresponding atomic coordinates for the TfuAlkX/DNA crystal structure have been deposited in the PDB under accession code 9ZD6 (https://www.rcsb.org/structure/9ZD6).

## Acknowledgements

This work was supported by grants from the National Science Foundation (MCB-1928918 to B.F.E.) and the National Institutes of Health (R01AI101171 and R01AI150701 to E.P.S.).

## Author Contributions

Y.C and D.E.K designed and performed experiments, analyzed data, constructed figures, and wrote the manuscript. M.E. performed experiments. B.F.E and E.P.S acquired funding, designed experiments, analyzed data, and edited the manuscript.

## Competing Interest Statement

The authors declare no conflict of interest.

## Notes

### Competing Interest Statement

The authors have declared no competing interest.

