## Supplementary for "DNA binding and lesion recognition by the bacterial interstrand DNA crosslink glycosylase AlkX"

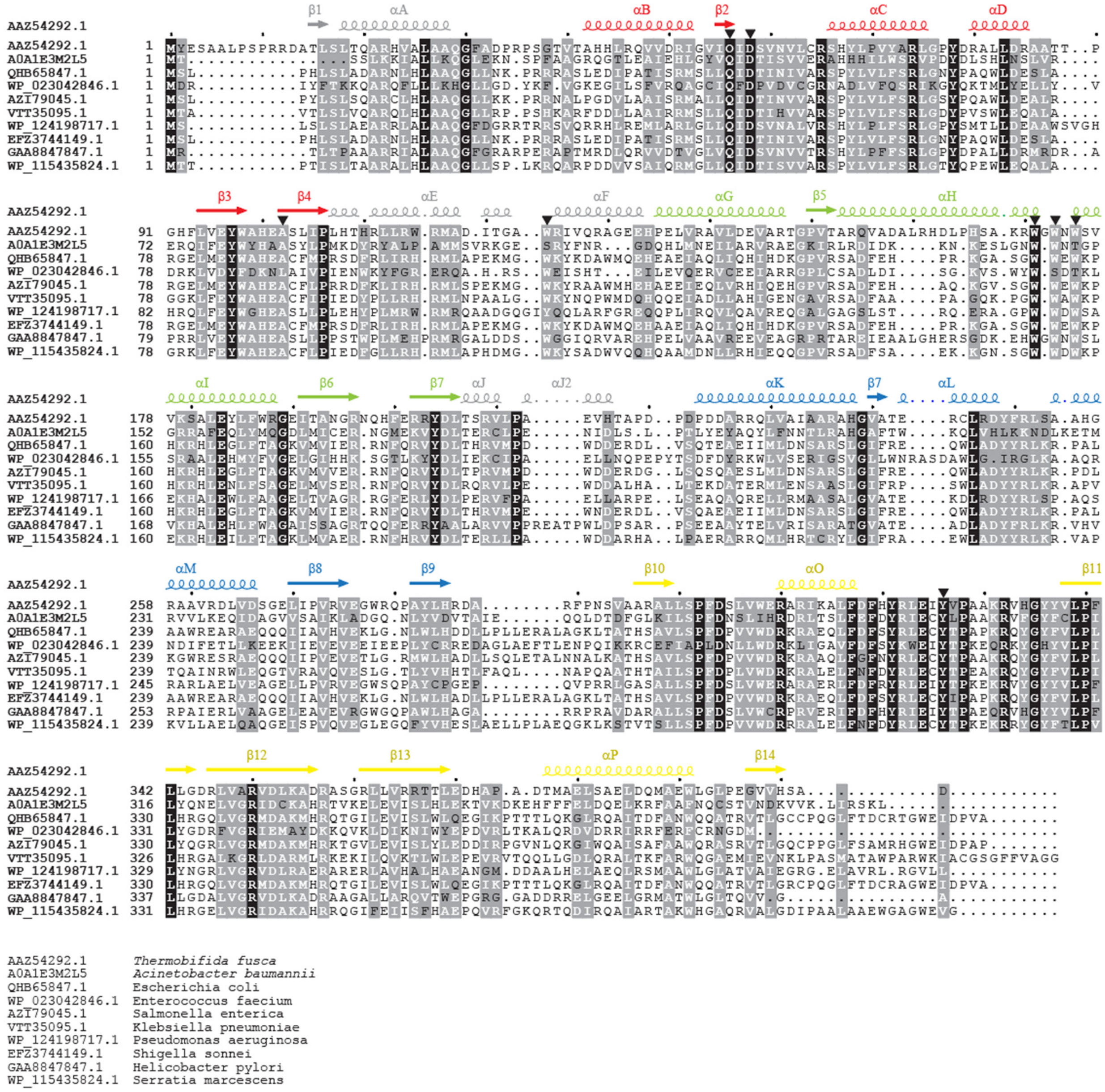

**Figure S1 Sequence alignment of YQL/AlkX proteins.** Secondary structure elements are colored by domain as in Fig. 1C. Black triangles denote residues mutated in this study.

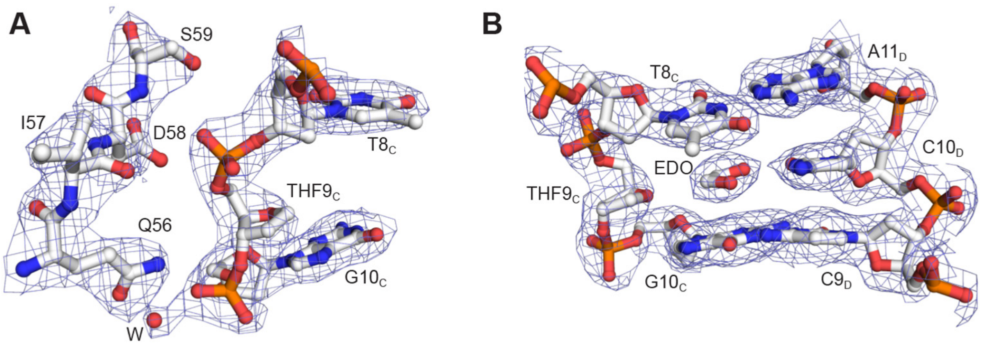

**Figure S2**. **TfuAlkX conservation.** The THF region of the DNA interacting with the active site (A) and the opposite DNA strand (B). 2Fo-Fc electron density contoured to 1σ is superimposed. Q56, I57, and D58 comprise the QxD catalytic motif.

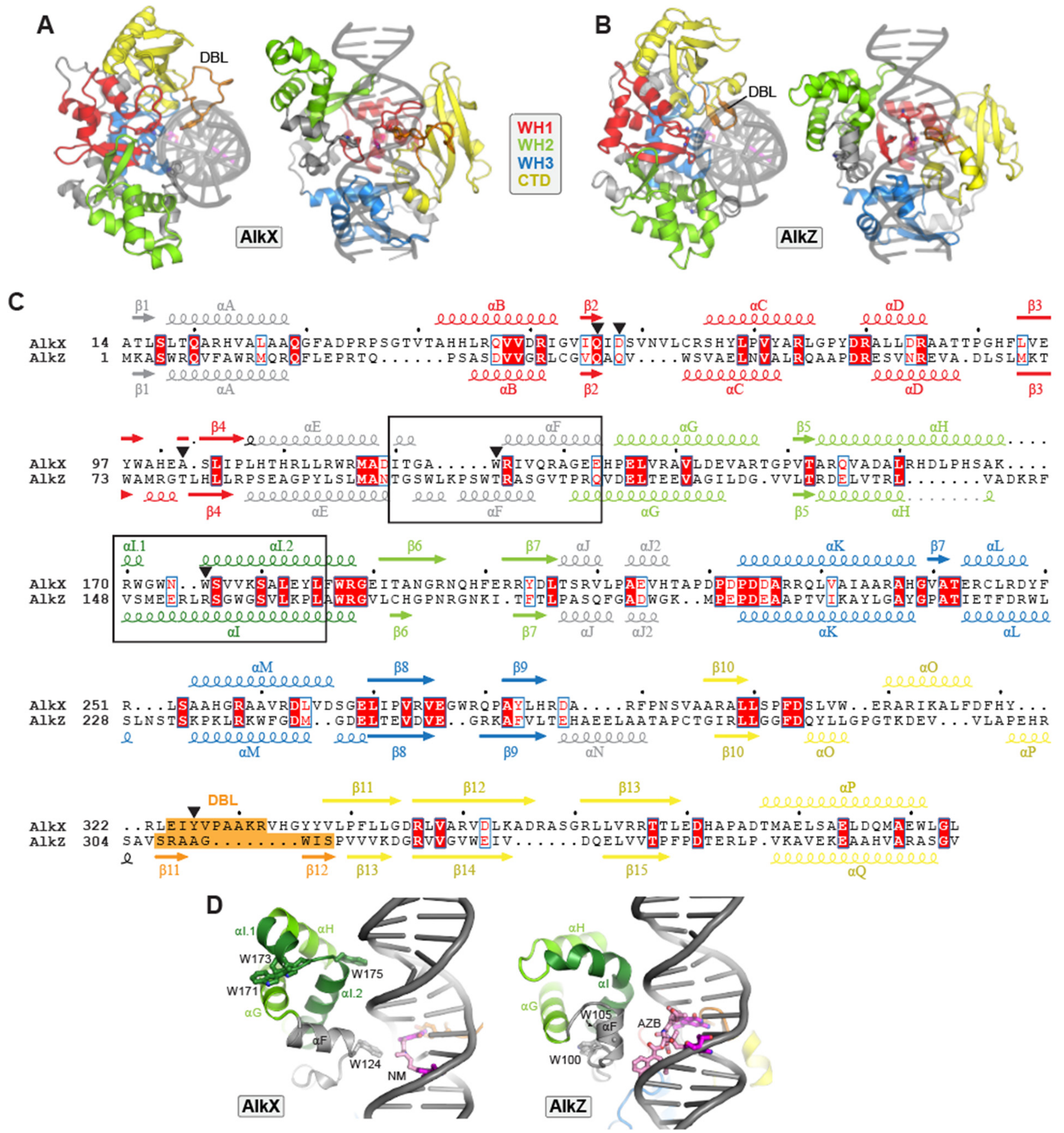

**Figure S3**. **Comparison of AlkX and AlkZ structures. A.** Crystal structure of TfuAlkX bound to THF-DNA. The protein is colored by domain (WH1 red, WH2 green, WH3 blue, C-terminal domain yellow, DBL orange). **B**. Crystal structure of *Streptomyces sahachiroi* (Ssa) AlkZ (PDB ID 5UUJ) docked against THF-DNA from the TfuAlkX structure. **C.** Sequence alignment of TfuAlkX and SsaAlkZ. Putative lesion sensing regions αF and αI are boxed. Residues important for ICL unhooking in AlkX are marked with black triangles. **D.** Comparison of αF and WH2 regions of AlkX (left) and AlkZ (right). DNA models of NM- and azinomycin B (AZB)-ICLs are based on the DNA in the TfuAlkX structure.

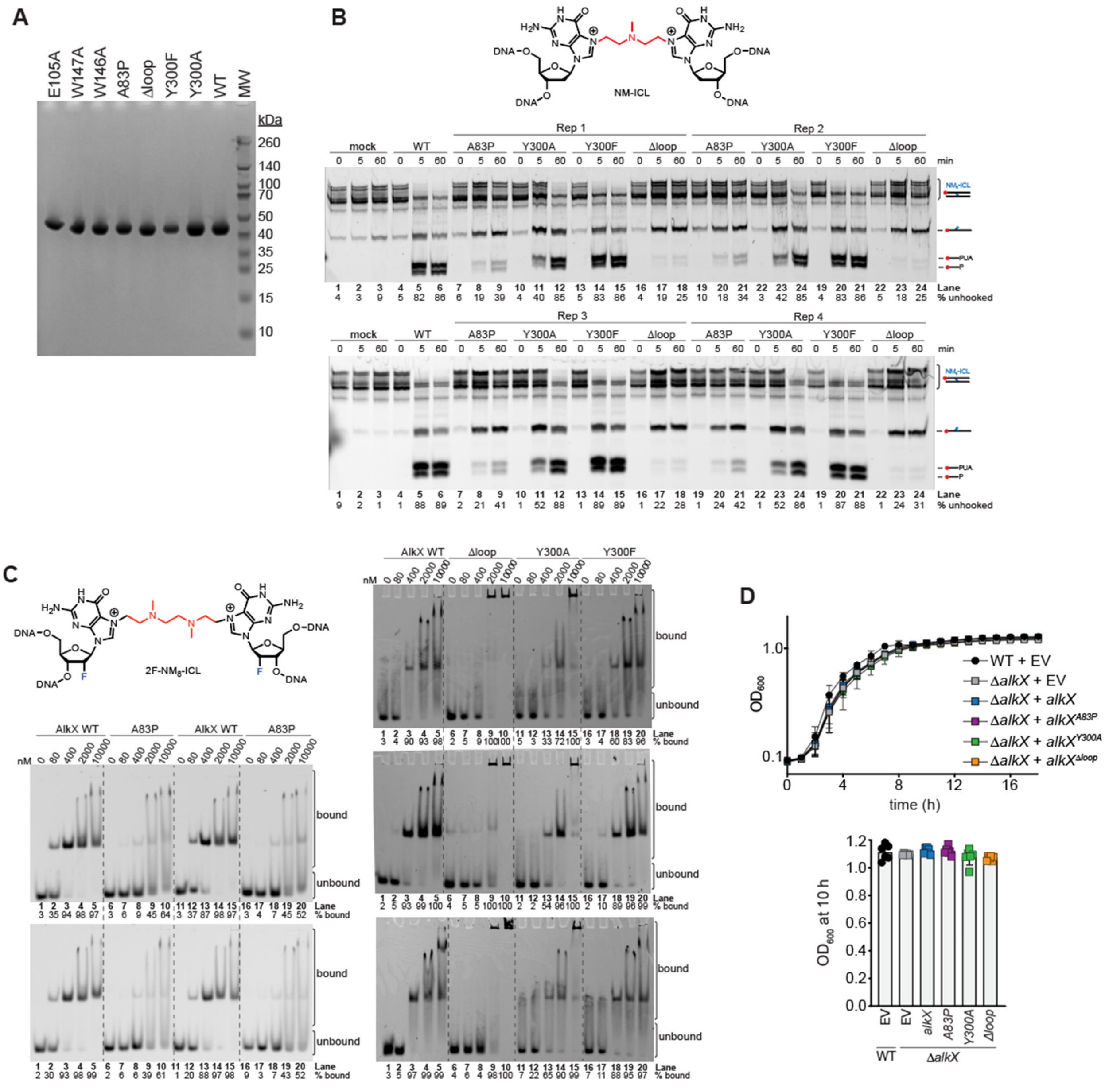

**Figure S4**. **Biochemical characterization of AbaAlkX mutants, related to data shown in Fig. 2 and Fig. 3. A**. SDS-PAGE of purified AbaAlkX proteins. **B**. The chemical structure of the NM-ICL used in base excision activity assays is shown at the top. Fluorescence (Cy5) scans of denaturing PAGE gels with four replicates of AbaAlkX reactions containing Cy5-labeled NM-ICL DNAat 0, 5 min, 60 min. Schematics to the right of the gel define the regions used in quantification. The AP-site product is cleaved during NaOH work-up to form a 3′-phospho-α,β-unsaturated aldehyde (PUA) and a 3′-phosphate (P). **C**. The chemical formula of 2F-$NM_8$-ICL DNA used in electrophoretic mobility shift assays (EMSAs). Fluorescence (Cy5) scans of native PAGE gels containing 3-4 replicates of AbaAlkX binding to Cy5-labeled 2F-$NM_8$-ICL DNA. **D**. AlkX complementation studies in untreated conditions. *Top*: Growth curves of *A. baumannii* strains (WT and Δ*alkX*) harboring pWH1266 empty vector (EV) or indicated pWH1266-*alkX* expression vectors in LB media. The $OD_{600}$ was recorded every 60 min. Data represent the mean ± SD of at least 6 biological replicates performed in technical triplicate. *Bottom*: $OD_{600}$ values at 10hr growth. Data represent the mean ± SD. Each dot represents an individual biological replicate, performed in technical triplicate.

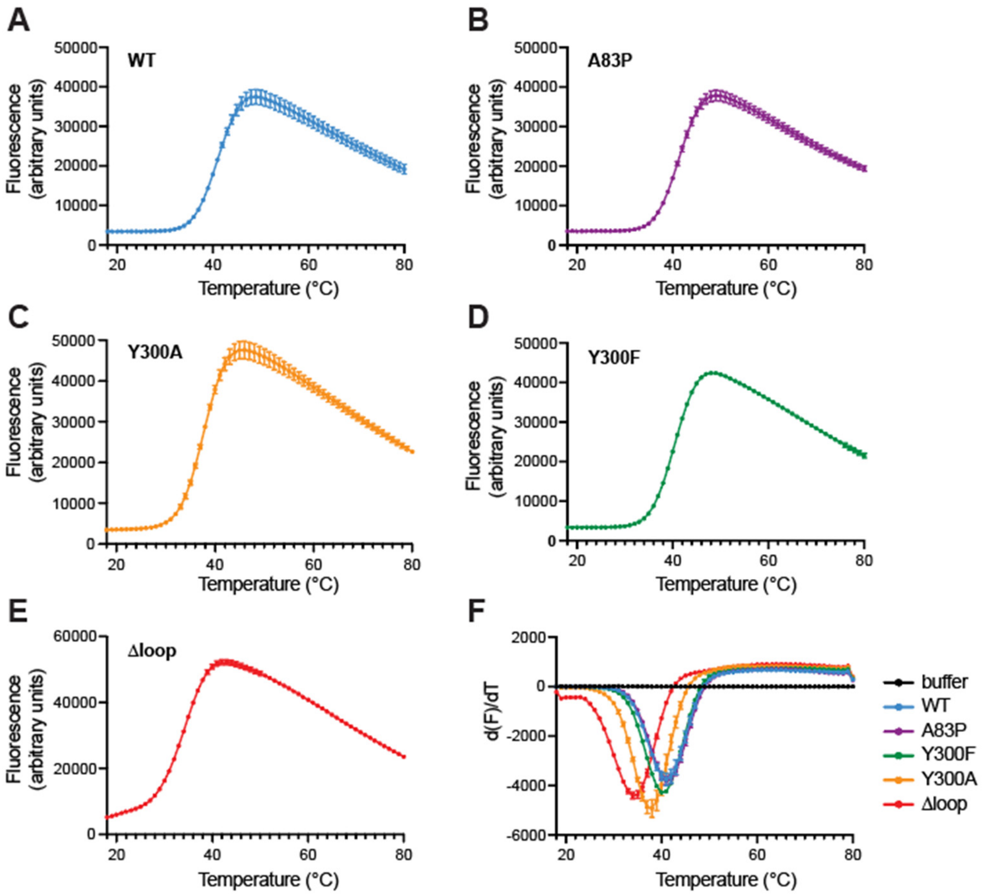

**Figure S5**. **Thermostability of AlkX mutants. A-E**. Differential scanning fluorescence thermal denaturation profiles for AlkX WT (A), A83P (B), Y300A (C), Y300F (D) and Δloop (E). Samples contained 10 µM protein, 20 mM Tris pH 8.0, 150 mM NaCl, 1 mM TCEP, 0.1 mM EDTA and 0.5X SYPRO Orange and measurements were carried out as previously described [Dorival and Eichman, *Nucleic Acids Res* **51**, 2838-2849 (2023). **F**. First derivatives of the thermal denaturation data in panels A-E. The minima signify the melting temperatures ($T_m$).

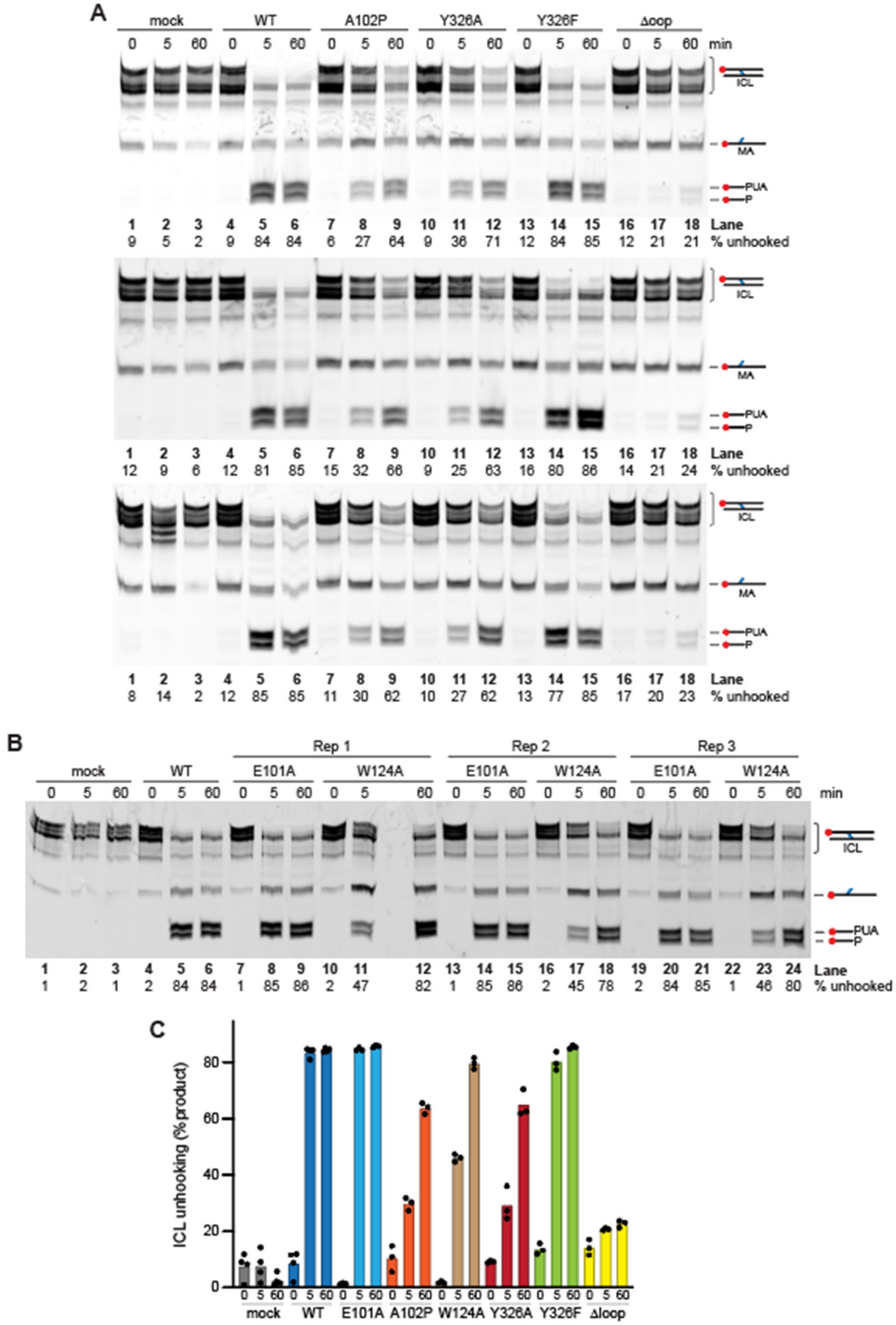

**Figure S6. ICL unhooking activity of TfuAlkX mutants**. **A,B**. Denaturing PAGE gels (Cy5 fluorescence scans) of reactions between TfuAlkX mutants and Cy5-labeled NM-ICL DNA, showing three biological replicates of A102P, Y326A, Y326F, Δloop (A) and E101A and W124A (B). **C**. Quantification of the gels. W124A data is related to data shown in Fig. 4.

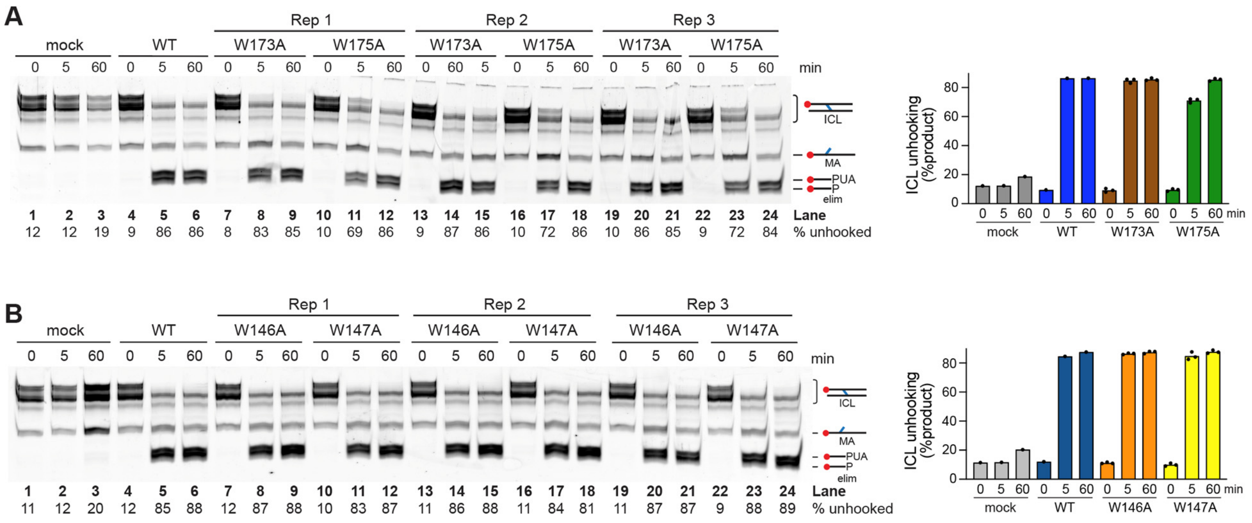

**Figure S7**. **ICL unhooking activity of residues in the αI region.** Denaturing PAGE gels of three biological replicates of NM-ICL unhooking reactions containing TfuAlkX (A) and AbaAlkX (B) mutants. Quantification of the gels is shown in the plots on the right.

**Table S1. X-ray data collection and refinement statistics**

| | |
|---|---|
| **Data collection** | |
| Space group | $P2_12_12_1$ |
| Unit cell | |
| *a, b, c* (Å) | 56.60, 115.64, 160.68 |
| *α, β, γ* (°) | 90, 90, 90 |
| Wavelength (Å) | 0.9655 |
| Resolution range (Å) | 56.60 - 2.60 (2.72 - 2.60)[a] |
| Total reflections | 214983 (25797) |
| Unique reflections | 33291 (4018) |
| $R_{merge}$ | 0.135 (0.983) |
| $R_{meas}$ | 0.147 (1.068) |
| $R_{pim}$ | 0.058 (0.413) |
| Mean I/σ(I) | 8.8 (2.3) |
| Completeness (%) | 99.9 (99.8) |
| Redundancy | 6.5 (6.4) |
| Wilson B-factor ($Å^2$) | 50.83 |
| $CC_{1/2}$ | 0.994 (0.801) |
| **Refinement** | |
| Resolution (Å) | 50.84 - 2.60 (2.69 - 2.60) |
| No. reflections | 33,197 (3,287) |
| $R_{work}$ | 0.2056 (0.3103) |
| $R_{free}$[b] | 0.2578 (0.3474) |
| No. atoms | 6,883 |
| Protein | 6,111 |
| DNA | 721 |
| Water | 46 |
| Other | 5 |
| Avg. B-factor ($Å^2$) | |
| Protein | 64.23 |
| DNA | 72.31 |
| Water | 55.54 |
| Other | 60.06 |
| Ramachandran distribution | |
| Favored (%) | 96.2 |
| Allowed (%) | 3.8 |
| Outliers (%) | 0 |
| RMS bonds (Å) | 0.009 |
| RMS angles (°) | 1.04 |

[a] Statistics for the highest resolution shell are shown in parentheses
[b] $R_{free}$ was determined from the 5% of reflections excluded from refinement

**Table S2. Strains, plasmids, and primers used in this study**

| Strain | Description | Reference |
|---|---|---|
| *E. coli* DH5α | Plasmid maintenance *E. coli* strain used for all cloning | Lab stock |
| *A. baumannii* 17978VU | Used for WT in this study | Lab stock |
| *A. baumannii* Δ*alkX* | 17978VU harboring an *ahp* kanamycin cassette insertion into the *alkX* locus | (1) |
| *E. coli* Tuner (DE3) | Protein expression | Lab stock |
| **Plasmid** | **Description** | **Reference** |
| pWH1266 | *Acinetobacter* expression plasmid | (2) |
| pWH1266-*alkX* | Expression plasmid encoding for *alkX* expression regulated by the native *alkX* promoter | (1) |
| pWH1266-*alkX*$^{A83P}$ | Plasmid encoding for the expression of a A83P mutant allele of *alkX*, regulated by the native *alkX* promoter | This study |
| pWH1266-*alkX*$^{Y300A}$ | Plasmid encoding for the expression of a Y300A mutant allele of *alkX*, regulated by the native *alkX* promoter | This study |
| pWH1266-*alkX*$^{\Delta loop}$ | Plasmid encoding for the expression of a Δloop mutant allele of *alkX*, regulated by the native *alkX* promoter | This study |
| pBG102 | pET27 derivative vector with N-terminal 6-his+SUMO tag for protein expression | Vanderbilt Center for Structural Biology |
| **Primer** | **Sequence** | |
| pWH1266_seq_F | TAGGCTTGGTTATGCCGGTACTG | |
| pWH1266_seq_R | GGAAGGAGCTGACTGGGTTGA | |
| pWH1266-AlkXA83P_F | GTATCATGCGccgTCTTATCTTCCtatgaaag | |
| pWH1266-AlkXA83P_R | caatattcgaaaatctgccgttcacgaaccaaac | |
| pWH1266-AlkXY300A_F | GATCGAATGCgcgTTACCAGCGGCcaaac | |
| pWH1266-AlkXY300A_R | cgataatcaaattcaaataaggaagttaaacggtc | |
| pWH1266-AlkX_Δloop_F | gcgaccacacccgtcctgtggatccTAAAGGTTTAGGTGAGTAAAG | |
| pWH1266-AlkX_Δloop_R | aaggctctcaagggcatcggtcgacTTAAAGCTTGCTGCGAATG | |
| AbaAlkXA83P_F | CATGCACCGAGCTATCTGCCGATGAAGGAC | |
| AbaAlkXA83P_R | CAGATAGCTCGGTGCATGATACCAGTACTC | |
| AbaAlkXY300A_F | GTATTGAGTGCGCTCTGCCAGCTGCCAAAC | |
| AbaAlkXY300A_R | CAGCTGGCAGAGCGCACTCAATACGATAGTC | |
| AbaAlkXY300F_F | GTATTGAGTGCtttCTGCCAGCTGCCAAAC | |
| AbaAlkXY300F_R | CAGCTGGCAGaaaGCACTCAATACGATAGTC | |
| AbaAlkXΔloop_F | CTATCGTATTGGCAGTAGTGGGGTCTTTGGCTACTTCTGCCTTCC | |
| AbaAlkXΔloop_R | GCCAAAGACCCCACTACTGCCAATACGATAGTCAAACTCAAACAAGC | |

| | |
|---|---|
| TfuAlkXA102P_F | GCACGAAccgAGCCTGATTCCGCTGCACAC |
| TfuAlkXA102P_R | cggAATCAGGCTcggTTCGTGCGCCCAATACTC |
| TfuAlkXY326A_F | CCGTCTGGAAATTgctGTGCCGGCGGCGAAAC |
| TfuAlkXY326A_R | GTTTCGCCGCCGGCACagcAATTTCCAGACG |
| TfuAlkXY326F_F | TCTGGAAATTtttGTGCCGGCGG |
| TfuAlkXY326F_R | CGGTAGTGAAAATCGAACAGC |
| TfuAlkXΔloop_F | GCTGTTCGATTTTCACTACCGTCTGggcagtagtgggGTGCACG |
| TfuAlkXΔloop_R | CGGCAGAACATAGTAACCGTGCACcccactactgccCAGACGGT |

[1] Kunkle DE, Cai Y, Eichman BF, Skaar EP (2024) An interstrand DNA crosslink glycosylase aids *Acinetobacter baumannii* pathogenesis. *Proc Natl Acad Sci U S A* 121: e2402422121

[2] Hunger M, Schmucker R, Kishan V, Hillen W (1990) Analysis and nucleotide sequence of an origin of DNA replication in Acinetobacter calcoaceticus and its use for Escherichia coli shuttle plasmids. *Gene* 87: 45–51
